# The longitudinal growth of the embryo of the kelp *Saccharina* depends on actin filaments that control the formation of an alginate corset in the cell wall

**DOI:** 10.1101/2024.07.10.603006

**Authors:** Samuel Boscq, Ioannis Theodorou, Roman Milstein, Aude Le Bail, Sabine Chenivesse, Bernard Billoud, Bénédicte Charrier

## Abstract

The initiation of embryogenesis in the kelp *Saccharina latissima* is accompanied by significant anisotropy in cell shape. Using monoclonal antibodies, we show that this anisotropy coincides with a spatio-temporal pattern of accumulation of alginates in the cell wall of the zygote and embryo. Alginates rich in guluronates as well as sulphated fucans show a homogeneous distribution in the embryo throughout Phase I of embryogenesis, but mannuronate alginates accumulate mainly on the sides of the zygote and embryo, disappearing as the embryo enlarges at the start of Phase II. This pattern depends on the presence of cortical actin filaments. In contrast, within the embryo lamina, the alginate composition of the walls newly formed by cytokinesis is not affected by the depolymerisation of actin filaments. Thus, in addition to revealing the existence of a mannuronate-rich alginate corset that may restrict the enlargement of the zygote and the embryo, thereby promoting the formation of the apico-basal growth axis, we demonstrate stage- and cytoskeleton-dependent differences in cell wall deposition in *Saccharina* embryos.

## Introduction

Brown algae are multicellular organisms that are characterised by a high diversity of body shapes (Charrier et al., 2012), ranging from microscopic uniseriate filaments (e.g. *Ectocarpus* spp; (Charrier et al., 2008; Le Bail et al., 2008)) to metres-long, thick blades (e.g. *Saccharina* spp; (Theodorou and Charrier, 2021). This great diversity of body shapes is due to the diversity of growth patterns. Brown algae feature different strategies for the spatio-temporal coordination of growth rate, growth location and growth direction in the three spatial axes, making them a model evolutionary group to investigate the various mechanisms underlying the acquisition of body axes and shape diversity in multicellular organisms.

The study of the embryogenesis of brown algae has led to the identification of several cellular and molecular factors inducing symmetry breaking. For example, the extremely polarised growth of the *Ectocarpus* filament depends on the Rho-GTPase pathway (Nehr et al., 2021), but no major triggering environmental factor has been identified. In contrast, the embryo of Fucales (e.g. *Fucus, Silvetia, Pelvetia* spp) polarises and germinates away from and parallel to the light source (Berger and Brownlee, 1994). The fertilisation site of the male gamete on the surface of the *Fucus* egg is also a positional cue triggering polarisation of the zygote (Hable and Kropf, 2000). This polarisation becomes apparent with the formation of a patch of actin filaments and the secretion of sulphated polysaccharides from the cell wall at the point where the rhizoid emerges (Bisgrove and Kropf, 2001; Hable et al., 2003). As with polarised cell growth in *Ectocarpus*, the transmission of these external signals involves the Rho-GTPAse pathway (Hable, 2014; Muzzy and Hable, 2013). To date, no studies of this type have yet been carried out on kelps (e.g. *Saccharina*). However, these brown algae have an interesting morphological feature. Unlike *Ectocarpus*, whose body plane is limited to the single axis of the uniserial filament (1D), and *Fucus*, whose embryo acquires its 3D architecture according to the sequence of the first three cell divisions of the thallus cell of the early embryo (Goodner and Quatrano, 1993), in Laminariales, embryogenesis is characterised by phases (or plateaus), during which growth is limited to first one axis (1D growth), then two axes (2D growth) before finally establishing the three perpendicular body axes making a 3D body (Kanda, 1941; Sauvageau, 1918). This feature makes them particularly amenable to the investigation of the factors controlling the establishment of body planes during embryogenesis.

We have recently described in detail these three morphological steps in the embryogenesis of *Saccharina,* particularly the transition from 2D to 3D growth, which, in this species, is accompanied by cell differentiation (Theodorou and Charrier, 2023). After fertilisation, the zygote first elongates along an axis parallel to the maternal stalk, which is the remnant envelope (cell wall) of the oogonium that sits on the gametophyte filament (Fig. 1A, left). The zygote thereby acquires its first body axis, which is the longitudinal axis (X-axis). Then, the second step corresponds to the maintenance of this initial body axis with a series of three cell division rounds perpendicular to the longitudinal axis of the zygote (each cell divides, almost synchronously) resulting in the formation of a stack of 8 cuboid cells (2^3^ cells, Fig. 1, left, Phase I; (Theodorou and Charrier, 2023). At this stage, the orientation of the next cell division tilts by 90°, which results in the formation of a second body axis, i.e. the medio-lateral axis (Y-axis) and the initiation of Phase II as defined in (Theodorou and Charrier, 2023) (Fig. 1, middle). After a series of alternations of transverse and longitudinal cell divisions, resulting in a monolayered lamina of ∼1000 cells (Fig. 1A, right), the cell division orientation again tilts by 90° in the third spatial axis (i.e. Z-axis), leading to lamina thickening (polystromatisation)– while continuing to grow in the other two axes – and cell differentiation.

**Figure 1:**
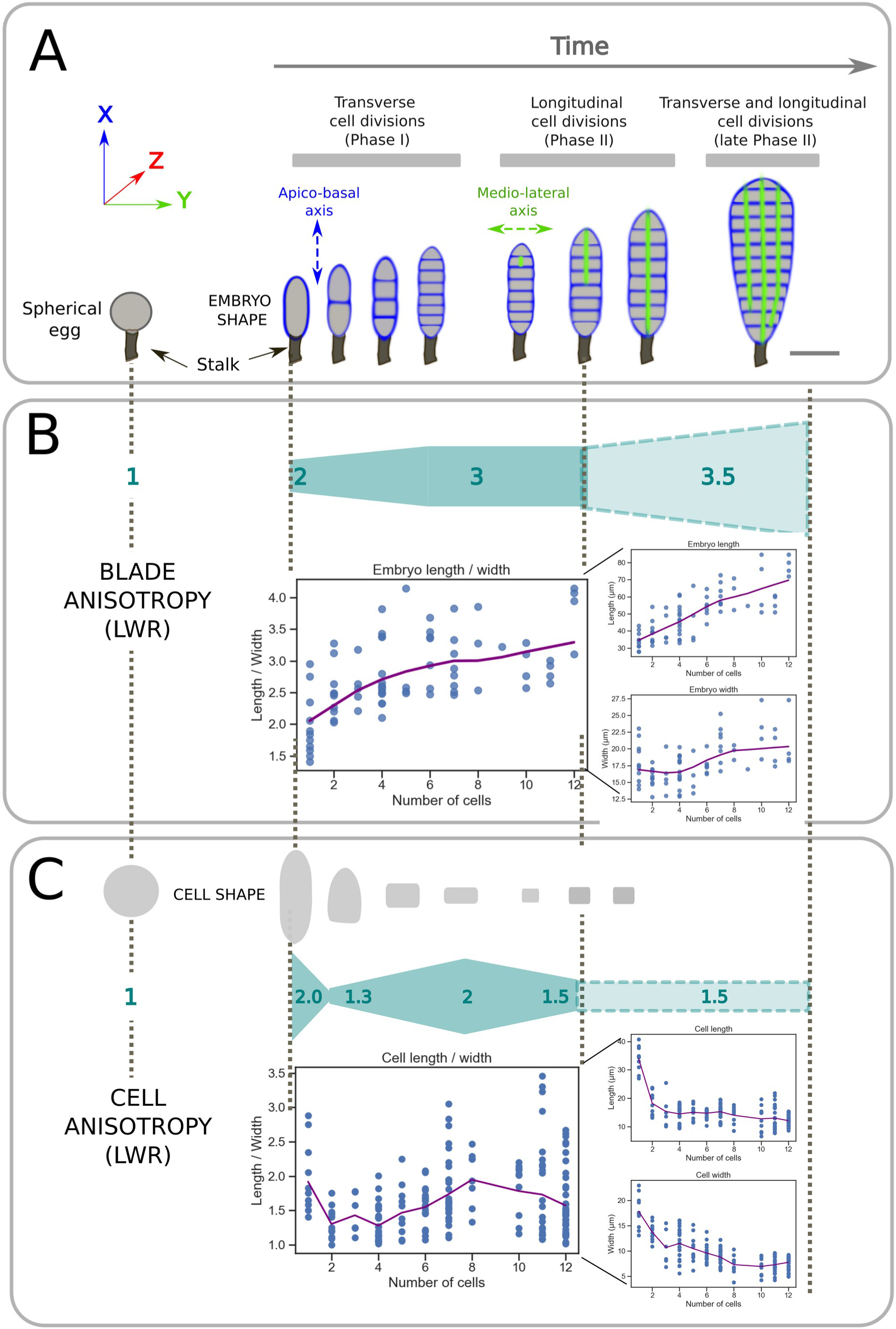
Anisotropy in the *Saccharina latissima* embryo. Development of the *S. latissima* embryo over time. (A) Schematics of the embryos from the zygote stage to Phase II (after cells divided longitudinally). Upon fertilisation, the zygote elongates (blue outline), establishing the first growth axis parallel to the maternal stalk (black peduncle) (left-hand side). Then, the early-stage embryo continues to develop longitudinally through transverse cell divisions (blue lines), forming the apico-basal axis (Phase I). Phase II starts with the first longitudinal cell division (green line), generally in the most apical region of the embryo (middle, top of the embryo). All cells eventually divide longitudinally, but this division is delayed in the basal cell. Scale bar = 50 µm. (B) Anisotropy in the morphology of the embryos. (Top) Schematised gradient of anisotropy along the development of the embryo, from the 1-cell stage (eggs, defined by LWR in [1.0;1.2] and zygotes), to the 12-cell stage (early Phase II). The numbers indicate the approximate value of the length-to-width ratio (LWR; unitless) at the corresponding stage. (Bottom) Graphics plotting the shape anisotropy value (expressed as the LWR, left), as well as the length (middle) and the width (right) over time (expressed as the number of cells of the embryo). The purple line shows the mean value, smoothed using the lowess function. (C) Anisotropy in the morphology of the embryo cells. (Top) Cell shape outlines at each developmental stage. Only recently divided cells are shown. (Middle) Schematised gradient of cell anisotropy along the development of the embryo. Cell LWR values are indicated. The purple line shows the mean value.

This exceptional series of ordered shifts, from 1D to 2D and then 3D growth at the level of the embryo, suggests that important positional cues are part of a regulation mechanism that promotes or suppresses the shifts in cell division orientations at one given stage. The maintenance of a given orientation of cell division over several division cycles is also an interesting feature, especially as in some cases it involves the formation of cells with a very high level of shape anisotropy, which, according to the canonical rules of cell division (Errera, 1888), requires specific constraints.

We have recently shown that the maintenance of the longitudinal axis depends on the contact of the embryo with the maternal stalk, which is the compartment from which the egg was extruded (Boscq et al., BioRXiv). This study suggested that a signal, most likely chemical, emanates from the maternal stalk (remnant oogonium), reaching and diffusing across the basal cell (Boscq et al., 2024a), and exerting acropetal control of longitudinal cell divisions. This control seems to be most prominent at Phase I during which the embryo reaches 8 cells, because severing the stalk before that stage results in embryos that grow isotropically.

At this stage of understanding the mechanisms controlling body axis establishment during the embryogenesis of *S. latissima*, we aimed first to precisely characterise the levels of anisotropy experienced by the embryo during its development, and second, to identify the molecular factors that might underlie this anisotropy.

## Results

### Embryogenesis in *Saccharina latissima* is accompanied by a complex pattern of embryo and cell morphological anisotropy

The sequential acquisition of the three body axes, characterised by plateaus during which the alga grows in certain axes, separated by abrupt transitions during which the alga starts growing in a new axis, involves 90° changes in the orientation of cell division at each transition. To determine whether the orientations of cell division depend on embryo and/or cell shape, we monitored several embryos during Phase I and early Phase II, and studied the relationship between emergence and maintenance of body axes in the embryo, and the degree of body and cell shape anisotropy.

After fertilisation, the spherical egg (length-to-width ratio (LWR)=1) elongates. This is the first anisotropy step of embryogenesis during which the embryo acquires it first body axis, which is the longitudinal apico-basal axis (Fig. 1A, left). As the embryo continues to grow, its shape becomes increasingly anisotropic (Fig. 1B; Table S1A) up to the 8-cell stage, where its LWR reaches 3.0, before dropping back to 2.5 at the 160-cell stage (Fig. S1A).

Detailed analysis of embryo length and width showed that, during Phase I, the embryo grew 36.2 µm in length while its width increased by only 2.7 µm (Fig. 1B, right; calculated from the 1-cell to the 8-cell stages; Table S1A). Thus, the anisotropy of growth of the embryo is due to a limitation of growth in width, i.e. along the medio-lateral axis (Y). Interestingly, width was stable at 17.6 µm until the ∼6-cell stage, when it started to increase. This corresponds to the transition from Phase I to Phase II, defined by the change in cell division orientation from exclusively transverse to both transverse and longitudinal. Therefore, the 90° tilt in cell division orientation coincides with an increase in embryo width.

The anisotropy dynamics of individual cells differ from that of the whole embryo. Once the zygote has divided, cells and the embryo have different trajectories of shape variation. As a whole, the anisotropy of the embryo continues to increase from the zygote to the 7-8 cell stage (Fig. 1B), but at the individual cell level, anisotropy varies more often (Fig. 1C; Table S1B): the zygotic cell had the highest degree of anisotropy (LWR≈2), after its elongation upon fertilisation. Then, due to the first embryonic cell division, cell LWR drops to 1.3, meaning that the first two embryonic cells are almost isotropic. The second and third rounds of transverse cell division increased the degree of anisotropy again to ≈2 (8-cell stage). This increase was due to the maintenance of transverse cell divisions with very little cell growth. As a result, not only does shape anisotropy increase — cells became much thinner in width (from 13.7 to 7.3 µm), while their length was almost unchanged (from 18.2 to 14.0 µm; Fig. 1C, right) —, but also the cell axis tilts by 90° (schematics in Fig. 1A, centre): initially aligned with the apico-basal axis of the embryo at the zygote and 2-cell stages, it aligned with the medio-lateral axis at the 4-cell stage onwards (Fig. 1C, top, centre). During the transitions from Phase I to Phase II, cell anisotropy began to decrease again, to LWR 1.5, which then remained stable throughout Phase II (Fig. S1C). In addition, at this stage, the cell axis alternated at almost each cell division.

Therefore, embryonic Phase I was characterised by two main events of morphological anisotropy: first, the elongation of the zygote, resulting from the retraction of the egg’s flanks after fertilisation, and second, recurrent transverse cell divisions that take place with virtually no cell growth and in the same orientation as the cell axis, resulting in highly anisotropic cells. The fact that this maximal morphological anisotropy was observed just before cells tilt their cell division planes suggests that high anisotropy would contribute to trigger the change in cell division orientation. Thus, to “bend the bow” (i.e. making cells highly anisotropic before tilting the cell division plane), there must be mechanisms that force cell divisions to be parallel to the longest cell axis.

### Alginate and fucan cell wall polysaccharides display different spatio-temporal patterns during embryogenesis in *Saccharina latissima*

To identify the cellular or molecular factors involved in the control of the complex anisotropy patterns characteristic of the *Saccharina* embryo, we first looked for potential molecular markers in the cell wall. The cell wall of brown algae is composed of mainly alginates and fucans that embed a small proportion of cellulose microfibrils (Charrier et al., 2019; Deniaud-Bouët et al., 2017; Mazéas et al., 2022). Chains of alginates are composed of different epimers, namely mannuronan acids (M) and guluronan acids (G) usually mixed within the chains as M-M, M-G or G-G blocks, which can be distinguished by the BAM6-11 series of monoclonal antibodies (Torode et al., 2016). Fucans can be present with different levels of sulphation, which can be targeted by the BAM1-4 series of antibodies (Torode et al., 2015). We took advantage of the specificity of these antibodies to try to identify molecular markers of anisotropy in the *Saccharina* embryo.

We observed that the eggs are enveloped by a homogeneous layer of mannuronate-rich alginates (i.e. rich in M-M blocks, labelled with monoclonal antibody BAM6) (Fig S2A, left). In contrast, in zygotes, they appeared to be present only along the flanks of the cell, with the apex and the base of the zygote being devoid of any signal (Fig. 2A). This corset of M-M-rich alginates surrounding the zygote persisted up to the 6-8 cell stage (Fig. 2B-G). Very interestingly, this corset disappeared as the embryo switched to Phase II, when transverse cell walls were labelled simultaneously (Fig. 2H-K). This pattern in which only the internal cell walls (both transversal and longitudinal), and not the external walls (corset) were labelled, persisted in Phase II embryos (Fig. 1L-0) and beyond (Fig S2A, right). It is noteworthy that M-M-rich alginates accumulated in the innermost layer of the cell wall of the cells within the embryo, so that two distinct cell walls were observed at each boundary between two cells, separated by a layer of cellulose microfibrils (Fig. 1P). Therefore, M-M rich alginates appear to be present in the most recent cell wall layers, which is in agreement with reports in *Fucus* embryos (Yonamine et al., 2021).

**Figure 2.**
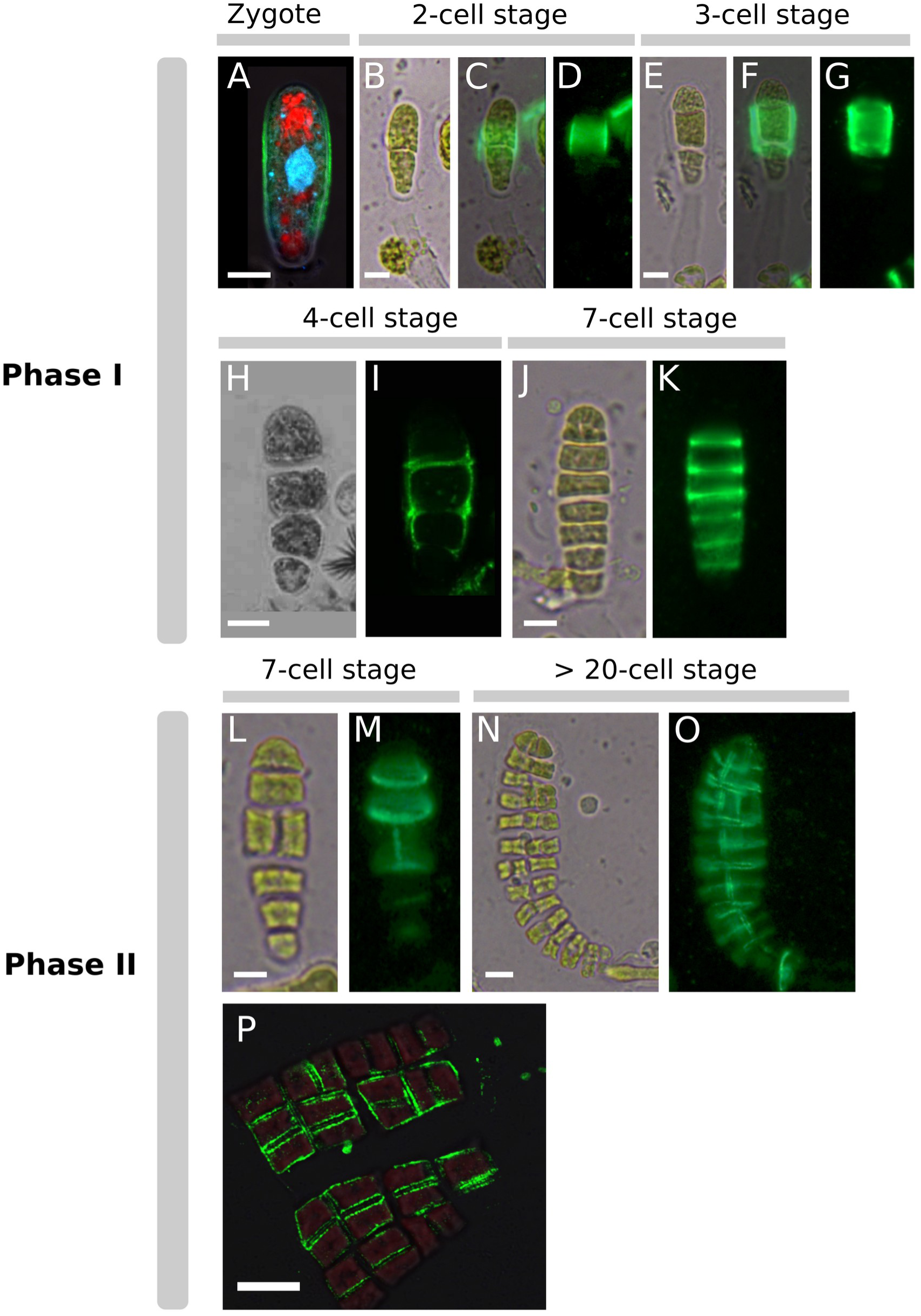
Detection of mannuronate-rich alginates during the development of *Saccharina* embryos. Zygote and embryos up to the >20-cell stages were labelled with the monoclonal antibody BAM6, which is specific to mannuronate-rich alginates. (A): zygote, (B-D): 2-cell stage embryo, (E-G): 3-cell stage embryo, (H,I): 4-cell stage embryo, (J,K): 7-cell stage embryo, (L,M): 7-cell stage embryo having divided longitudinally (formally, a Phase II embryo), (N,P): Phase II embryos (with transverse and longitudinal cell division. (B,E,H,J,L,N): Bright field images. (D,G,K,M,O): epifluorescence of FITC-BAM6 antibody (green signal). (C,F): merge. (A,H,I,P): confocal images. (P):close-up of a blade of a Phase II embryo showing mannuronate-rich alginates in the innermost cell wall of each cell (green: FITC; red: chloroplast autofluorescence; blue: Calcofluor; cyan: DAPI). Scale bars = 10 µm.

In summary, M-M-rich alginates were found in all cells, except in the apex of embryos. From our observations, in the early stages, S*accharina* embryos do not grow apically; instead growth is diffuse as all cells grow. Therefore, the absence of M-M-rich alginates in the apex raises questions about their role in *Saccharina* embryogenesis.

The BAM7 antibody, which labels M-G-rich alginates (Torode et al., 2016), did not display the same spatio-temporal pattern as the BAM6 antibody (Fig. S2B). In contrast to M-M-rich alginates, M-G-rich alginates were homogeneously distributed in fertilised eggs and zygotes, not only along the flanks, but also at the apex and the base of the embryo. In addition, right after the zygote stage, these alginates were no longer detected in the external cell wall and accumulated in the internal cell walls only (Fig. S2B). At the sub-cellular level, M-G-rich alginates were detected in the innermost layer, like M-M-rich alginates (Fig. S2B).

Guluronate- (G-G-)rich alginates, labelled by monoclonal antibody BAM10 (Torode et al., 2016), were detected at the surface of embryos from the egg stage to mature Phase II (Fig. S2C, right). In comparison, these polysaccharides were barely detected in internal walls, where the signal was much weaker than that of Calcofluor white, which binds to cellulose microfibrils (Fig. S2C, blue channel).

A similar pattern was found for fucans, which are abundant components of cell walls in most brown algae (Charrier et al., 2019; Popper et al., 2011). Using the monoclonal antibody BAM4 (Torode et al., 2015), we observed that sulphated fucans uniformly envelop the surface of eggs, zygotes and embryos until late Phase II at least (Fig. S2D), but not in the internal cell walls of Phase II embryos (Fig. S2D). Interestingly, like G-G-rich alginates in the enveloppe of eggs and zygotes (Fig. S2C), sulphated fucans were detected more externally than cellulose (in e.g. zygote in Fig. S2C and Phase I and II embryos in Fig. S2D).

In contrast to alginates and fucans that are present in specific locations during the development of *Saccharina* embryo, cellulose labelled with Calcofluor white appeared to be distributed homogeneously in the external and in all the internal cell walls from the egg to the Phase II stages (Fig. S2A-D).

### Actin filaments are localised cortically in Phase I and Phase II embryos

In some brown algae belonging to the orders Fucales and Sphacelariales, the accumulation of cell wall components depends on the organisation of actin filaments (AFs) (Hable, 2014; Hable and Kropf, 2005; Hable et al., 2003; Hable et al., 2008; Henry et al., 1996; Katsaros et al., 2003; Muzzy and Hable, 2013). To assess the role of AFs in the formation of the cell wall during *Saccharina* embryogenesis, and more specifically in the establishment of its morphological and molecular anisotropy, we labelled AF-fixed embryos with rhodamine-phalloidin or Alexa Fluor^TM^ 488 phalloidin. In zygotes, we observed a network of cortical AFs throughout newly formed zygotes (Fig. 3A,B, white arrows). In very anisotropic zygotes, the stains were sometimes concentrated mainly along the flanks (slightly stronger signal in Fig. 3C,D; white arrow). This accumulation in the cell cortex was maintained in Phase I embryos (Fig. 3E,F), and even more clearly in Phase II, where all the innermost sides of the cuboid cells that characterise Phase II were homogeneously labelled (Fig. 3G-J). The cortical localisation of AFs is not specific to the *Saccharina* embryo, having been observed in other brown algae, particularly when cells are in interphase (Katsaros et al., 2006). Interestingly, a weaker signal was observed at the periphery of the embryo blade, along the walls on the outward-facing edges of the embryo (Fig. 3H,J, white asterisks). This distinct pattern suggests that the role of AFs in the cells of the lamina differs between the inner edges shared by neighbouring cells and the outer edges where cell faces are in direct contact with the seawater. This interpretation is supported by the different orientations of AFs in Phase II embryos (Fig. S3), which are all aligned along the Z-axis perpendicularly to the embryo apico-basal axis (Fig. S3A,C; angle= 77.69° ± 10.43; Fig. S3D, left; Fig. S3E), except when they are close to the front and back of the embryos, along the cell faces in contact with seawater (Fig. S3B; angle= 50.37° ± 26.41; Fig. S3D, right). This highly ordered organisation may be involved in the differential organisation of the cell wall that underlies the anisotropy of cell wall expansion.

**Figure 3.**
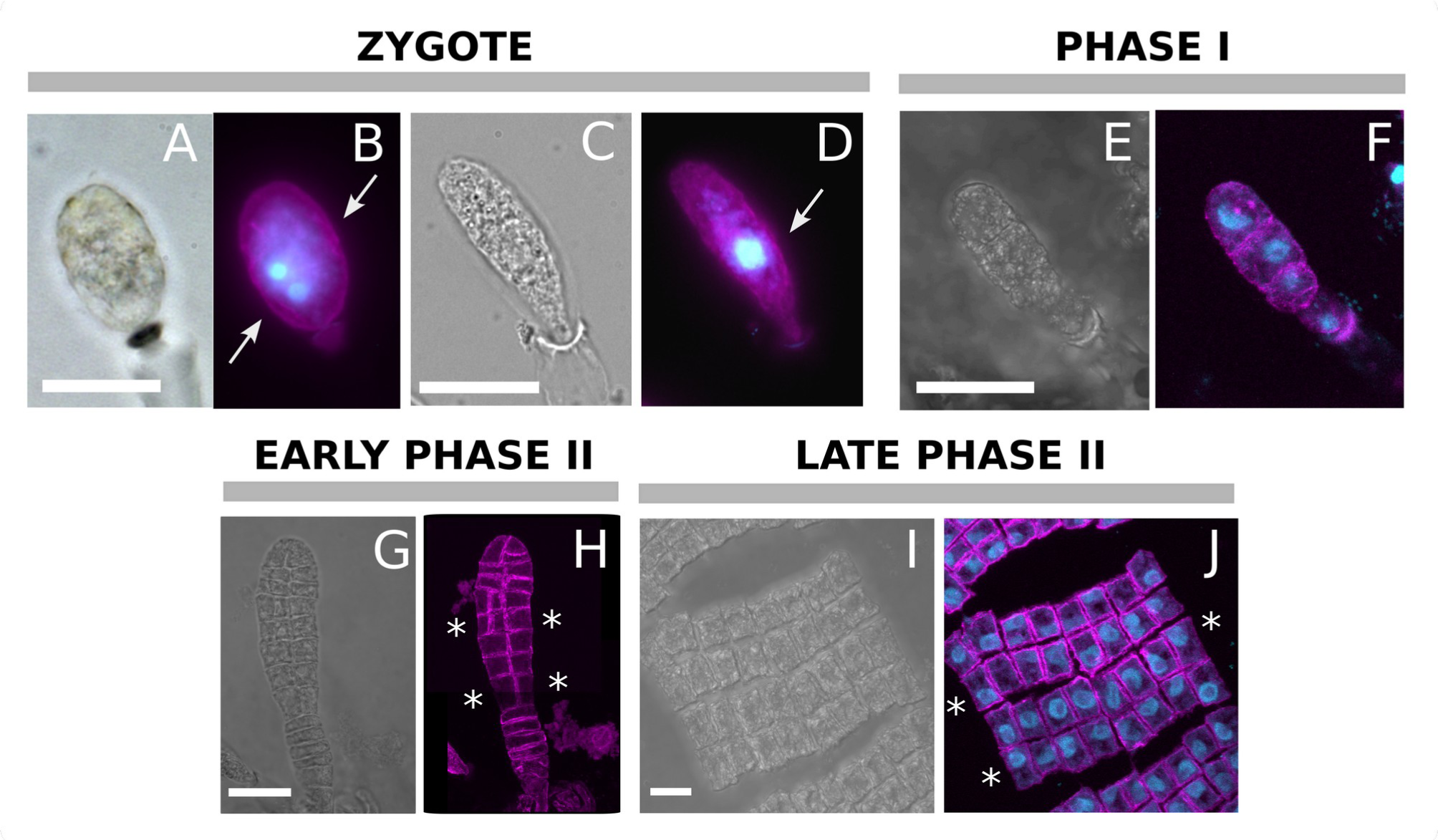
Actin filament organisation during *Saccharina* embryogenesis. (A-D): zygotes, (E,F): Phase I embryos, (G,H): early Phase II embryos, (I,J): late Phase II embryos. (A,C): transmission light images of epifluorescence microscopy of specimen shown in B and D; (E,G,I): transmission light images from confocal microscopy of specimens F, H and J, respectively. (B, D): confocal image (single optical section); (F, H, J): maximal projections of confocal Z-stacks. Actin filaments (AF, magenta) in B, D and J are stained with Alexa Fluor^TM^ 488-phalloidin conjugate and in F and H with rhodamine-phalloidin. Nuclei (cyan) were stained with DAPI in images B, D, F and J. They display an apparently random position within each cell. White arrows indicate cortical labelling. White asterisks indicate the weaker signals on the outer edges of the embryo. Scale bars = 20 μm.

### Actin filaments control cell shape

To further characterise the process that controls the specific organisation of the cell wall in *Saccharina*, we treated the *Saccharina* embryo with latrunculin B (LatB), a drug that binds actin monomers and blocks their polymerisation into AFs (Wakatsuki et al., 2001). Embryos were treated for 7 days with 50, 100, 500 or 1000 nM LatB in seawater. Control embryos immersed in 1% DMSO grew with no malformation (Fig. 4A), but LatB-treated embryos showed reduced growth and displayed abnormal morphologies (Fig. 4B-I). These morphological responses were observed for LatB concentrations as low as 50 nM (Fig. 4B,C), for which eggs and zygotes grew into cells larger than the control, and with irregular shapes. The extent of the impact of LatB on the morphology of the embryo increased with LatB concentrations from 100 nM (Fig. 4D) up to 1 µM (Fig. 4E-I). Zygotes swelled and grew asymmetrically, generating protuberances in all three spatial directions (e.g. Fig. 4B), or a pair of spherical cells parallel to each other in the medio-lateral direction (Fig. 4G). Some returned to an isotropic shape, losing their elongated shape and apico-basal axis (Fig. 4E,I). These abnormal morphologies indicate that AFs are involved in maintaining the elongated shape of the zygote, and in controlling the orientation of divisions during Phase I, when it occurs in these conditions (AFs are also necessary for cytokinesis (Katsaros et al., 2006), and LatB treatment usually inhibits cell division). However, the effect of LatB on the orientation of cell divisions diminished once the first transverse division occurred, and only cell shape remained affected (Fig. 4D,H). Eggs, whether recently fertilised or not (spherical cell), may adopt an irregular shape (Fig. 4C).

**Figure 4.**
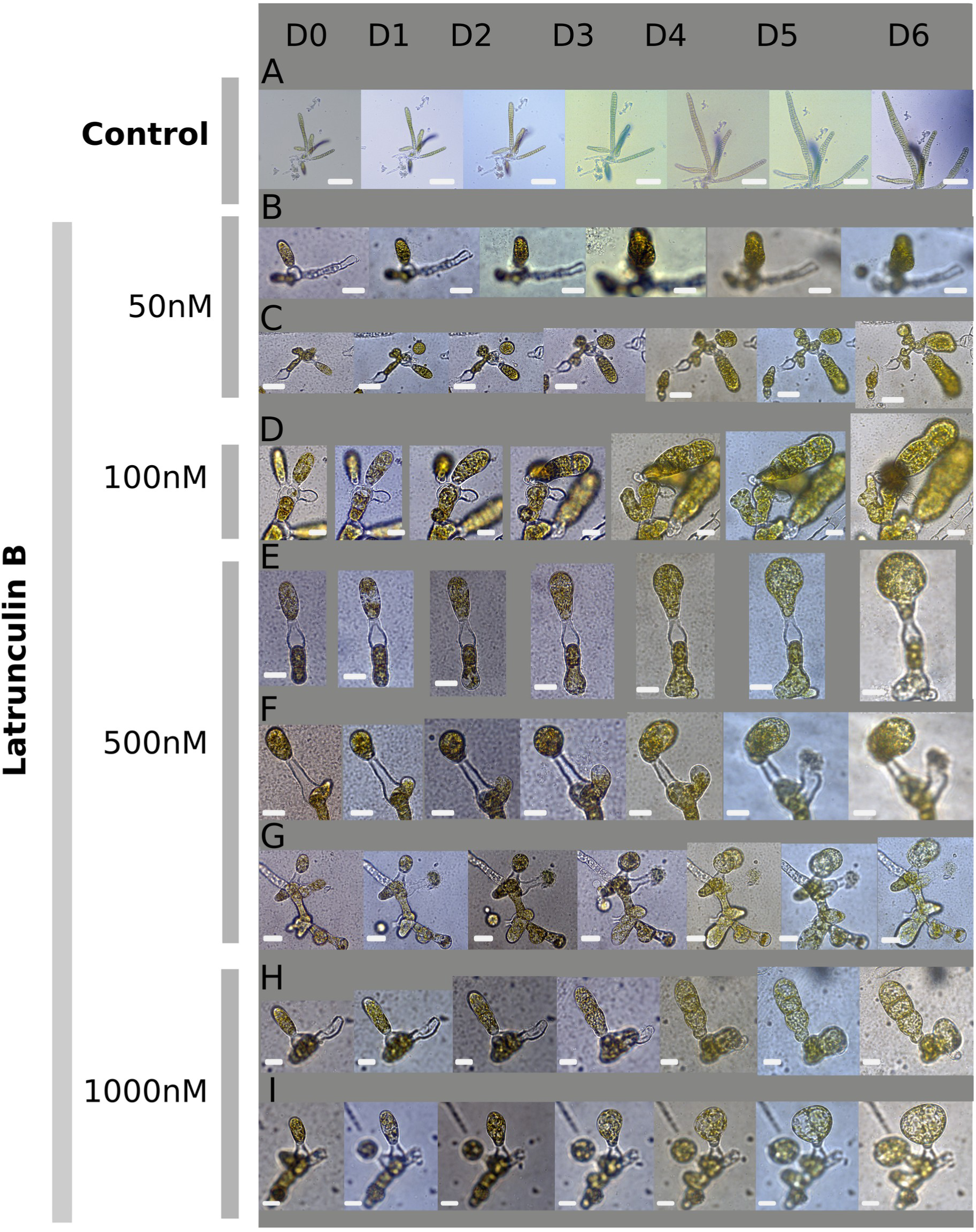
Morphological response of *Saccharina* zygotes treated with Latrunculin B. Zygotes (D0) were treated with increasing latrunculin B concentrations for 7 days (D0 to D6). A photo is shown for each day for treated zygotes (B-I) and for DMSO controls (A). (B,C): 50 nM, (D) 100 nM, (E-G): 500 nM; (H,I): 1 µM. Scale bars = (A) 200 µm, (B,C) 20 µm, (D-I) 10 µm.

### Actin filaments control the spatio-temporal pattern of mannuronate-rich alginates during *Saccharina* embryogenesis Phase I and Phase II

To assess whether AFs are functionally related to the composition of the cell wall, and specifically involved in the anisotropic pattern of M-M-rich alginates along the flanks of the embryos, we repeated the labelling experiments of alginates and fucans on eggs, zygotes and embryos in the presence of LatB. When treated with 1 µM LatB for 1 week, eggs, zygotes and embryos showed reduced anisotropy, as observed in Fig. 4. In addition, M-M-rich alginates accumulated in the apex of zygotes in addition to the flanks (Fig. 5A-F), unlike control embryos (Fig. 2 and text above). Labelled M-M-rich alginates in the tip of the embryos persisted to the 7-cell stage (Fig. 5G-J): the first apical half of the embryos was homogeneously labelled in addition to the transverse cell walls (also Fig. S2A). Therefore, in response to LatB, the “corset” of M-M-rich alginates, limited to the flanks of the control embryos, is transformed into a “sock” covering both the flanks and the apex.

**Figure 5.**
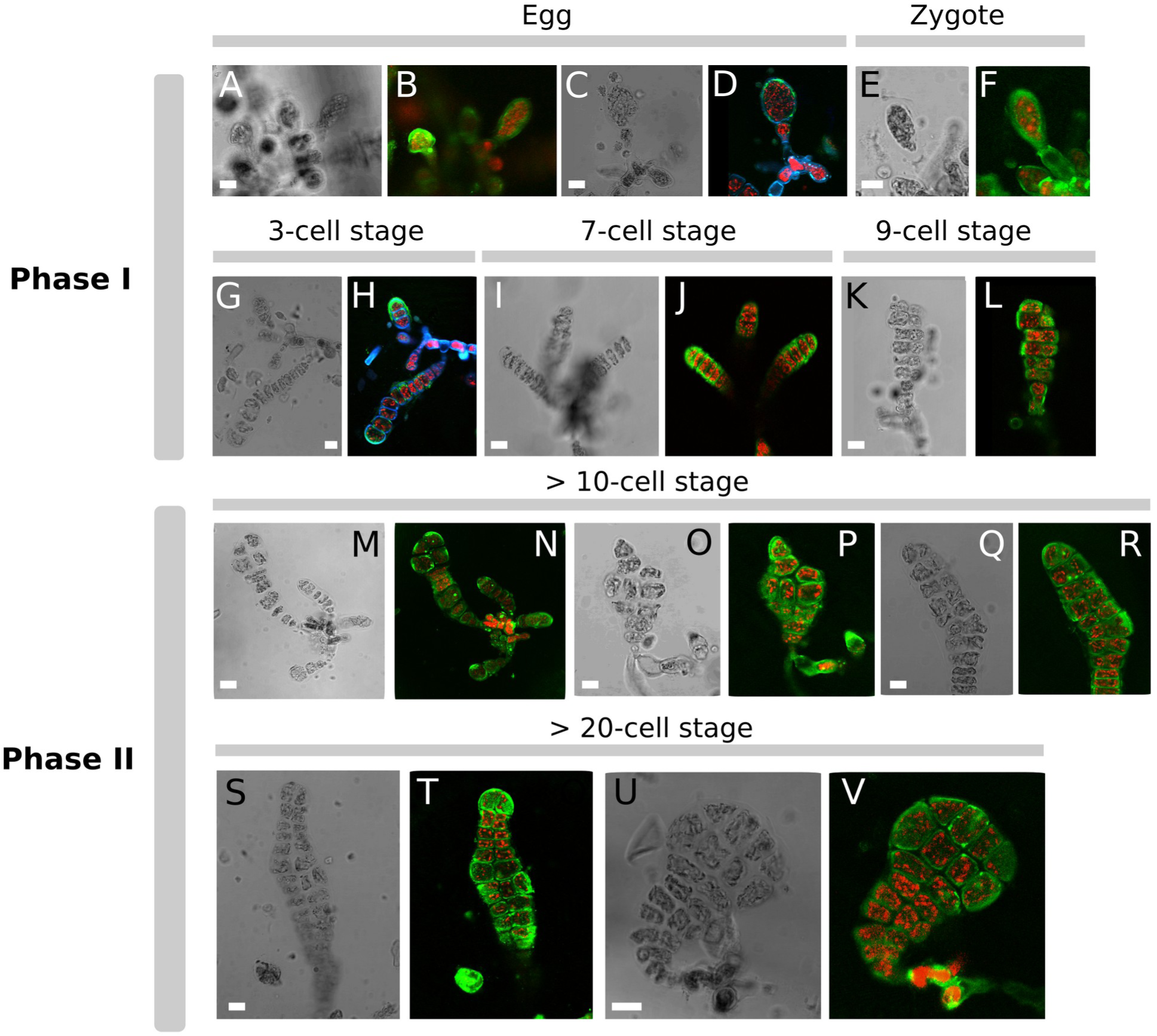
Labelling of mannuronate-rich alginates in *Saccharina* embryos treated with latrunculin B. Fertilised eggs, zygotes and embryos were labelled with the monoclonal antibody BAM6 after having been treated by 1 µM latrunculin B for 7 days. (A,C,E,G,I,K,M,O,Q,S,U): bright field images. (B,D,F,H,J,L,N,P,R,T,V): confocal images of FITC-BAM6 labelling (green: FITC; red: chloroplast autofluorescence; blue: Calcofluor white). Scale bars = 10 µm.

In Phase II embryos, following longitudinal cell divisions, M-M-rich alginates were observed on the boundary of the blade (Fig. 5M-V; Fig. S4A). This pattern contrasts with the pattern observed in control embryos in which the blade boundary was devoid of M-M-rich alginates. The labelling of transverse cell walls in response to LatB remained unchanged (Fig. 5R,T,V). In summary, LatB-mediated depolymerisation of AFs resulted in a ubiquitous accumulation of the BAM6 epitope in the periphery (outer cell wall) of the egg, zygote and embryos and in the transverse cell walls within the lamina. This ubiquitous accumulation of M-M rich alginates, maintained from the egg stage until the end of Phase II, is in contrast to the change observed between Phase I and Phase II in control embryos.

Thus, AFs play a role in the control of mannuronate-rich alginate accumulation or organisation in the external cell wall that grows and thickens as of the zygote stage, but not in the transverse and longitudinal internal cell walls that are formed by cytokinesis.

Remarkably, this regulation of the spatio-temporal accumulation pattern is highly specific to M-M-rich alginates, because no such modification was observed in the presence LatB when using BAM7, and BAM10 antibodies (compare Fig. S4 with Fig. S2). However, it is noteworthy that the alginate signal was less well defined in the presence of LatB than in control growth conditions. Uneven patches of FITC were observed when using BAM6 and BAM7 (Fig. S4C), suggesting a major misregulation of M-M-rich and M-G-rich alginate accumulation in these embryos. Altogether, this result supports that the localisation of M-M-rich alginates, and to a lesser extent M-G-rich alginates, is impaired in LatB-treated *Saccharina* embryos.

Interestingly, LatB did not affect the localisation of cellulose and sulphated fucans, because these cell wall polysaccharides displayed a similar distribution as that described in control conditions (Fig. S2). In the presence of LatB, sulphated fucans still enveloped the zygotes and Phase I and Phase II embryos (Fig. S4D), similarly to G-G-rich alginates. Similarly, cellulose microfibrils were present in the external and internal cell walls in the same manner as in the control conditions (easily observed in Fig. S4D for example, Phase I embryos). Therefore, in the early development of *Saccharina* embryo, neither the location of sulphated fucans, nor the formation of cellulose microfibrils depend on AF networks.

## Discussion

In this study, we aimed to test whether the assembly of cell wall polysaccharides work in concert with cortical actin filaments to contribute to the dynamics of anisotropic growth of the *Saccharina* embryo, principally during Phase I when the first body plane, namely the apico-basal axis, is established. First, we immunolocalised alginates and fucans in the cell wall of *Saccharina* eggs, zygotes and embryos. We showed that, in contrast to the other cell wall polysaccharides tested in this study, the accumulation pattern of mannuronate-rich alginates, coincides in space and time with the establishment of the longitudinal axis of the zygote (summarised in Fig. 6, top). We also showed that AFs are necessary for the zygote to establish or maintain this longitudinal axis (Fig. 6, bottom). These results prompt the following question: can mannuronate-rich alginates regulate the formation of the longitudinal axis and, if so, how?

**Figure 6.**
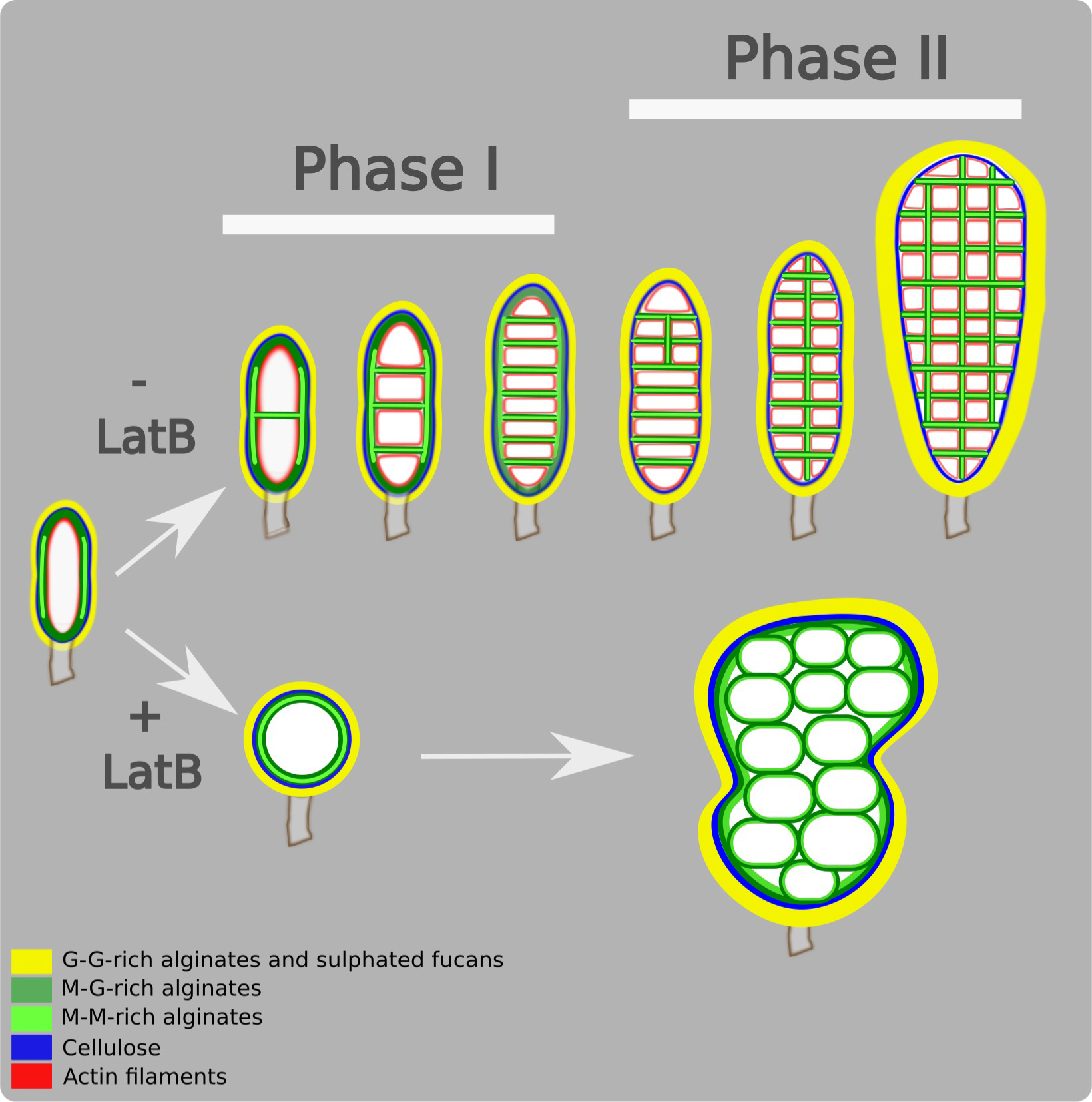
Summary schematics of the localisation pattern of the cell wall polysaccharides in zygotes and Phase II embryos. Schematics representing the typical body shape and the localisation of cell wall polysaccharides in the zygote, Phase I and Phase II embryos of *Saccharina latissima*, in standard culture conditions (top) or in the presence of latrunculin B (LatB; bottom). In LatB-treated material, the zygote is spherical and the embryo deformed, made of rounder cells. A layer of mannuronate-rich alginates homogeneously envelops the whole zygote and persists up to late Phase II when the embryo is made of several rows and columns of cells, which may be misaligned. For better clarity, cellulose (in blue) is not represented in the inner cell walls. M, mannuronate residue; G, guluronate residue.

There are many explanations for changes in the immunochemical labelling pattern. This change can be attributed to processes such as degradation, changes in conformation, synthesis or delivery of another compound obscuring the targeted epitope (Hawes et al., 2009; Vázquez-Gutiérrez and Langton, 2015). Whatever the reasons for the signals appearing and disappearing, our experiments show that polysaccharides fall into two categories based on the spatio-temporal occurrence in the cell walls of the *Saccharina* embryo. The first category, made of guluronate-rich alginates and sulphated fucans, was observed in outer cell walls only (Fig. 6, top). In this compartment, these polysaccharides were homogeneously distributed in the most external layer of the cell wall of the whole embryo. Their localisation pattern did not change from the egg stage to the Phase II embryo. These polysaccharides are a permanent component of the thick, external cell wall enveloping the embryo as of the egg stage and are weakly detected in the internal cell walls. Many brown algal embryos synthesise an outer envelope very quickly after fertilisation. In *Saccharina*, the cell wall forms quickly after fertilisation ((Motomura, 1990); and our observation). In Fucales, the zygote starts forming a fibrous cell wall less than 5 minutes after fertilisation (e.g. *Fucus vesiculosus,* (Pollock, 1970), which is surrounded by a multilayered cell wall ∼500 nm thick 4-6 h after fertilisation, and *Pelvetia compressa*, (Bisgrove and Kropf, 2001). Similarly, in Dictyotales, cell wall secretion is detected within 90 s after fertilisation (e.g. *Dictyota dichotoma* (Bogaert et al., 2017b). Thus, our results are consistent with this general pattern.

The second category of cell wall polysaccharides was observed in specific locations, which varied during the growth of the embryo. M-M-rich and M-G-rich alginates were first localised on the surface of the zygote, and then in the transverse and longitudinal internal cell walls of Phase I and Phase II embryos (summarised in Fig. 6, top). Therefore, their spatial pattern changed with time. Furthermore, and very interestingly, M-M-rich alginates were not detected in the apex of zygotes and Phase I embryos, in contrast to M-G-rich alginates. Instead, they formed a corset on the sides of the zygote and early embryos (up to the 8-cell stage) and, remarkably, this corset disappeared when the embryo initiated growth along the second body plane: the medio-lateral axis. Upon fertilisation, the egg elongates and preliminary data suggest that the elongation of the *Saccharina* egg is not accompanied by an increase in cell volume. This elongation without enlargement implies that there must be a mechanism to retract the flanks inwards along the Y and Z axes for the egg to increase in length along the longitudinal axis (X-axis). This mechanism has also been described in the zygote of *Dictyota*, which elongates less than 90 s after fertilisation through a mechanism of actomyosin-mediated cell contraction resulting in the modification of cell shape without growth (Bogaert et al., 2017a). In addition, in *Saccharina latissima*, longitudinal cell divisions do not occur until the embryo reaches ∼8 cells, which corresponds to Phase I (Theodorou and Charrier, 2023). Therefore, during Phase I, growth in width is limited in terms of cell expansion and cell division (increase in width =16.7%) while the embryos double in length (increase of 114.1%). Thus, the corset of mannuronate-rich alginates, which persists until the 8-cell stage in Phase I embryos, may ensure that cells do not expand laterally by stiffening the cell wall. At the end of Phase I, the corset disappears and the cells begin to divide longitudinally, allowing the embryo to widen. Knowing whether alginate epimers also accumulate anisotropically along the longitudinal axis of the zygote of *Dictyota,* at the surface of which the cellulose layer seems homogeneously distributed (Bogaert et al., 2017b), would open the way for comparative studies on the role of mannuronate alginates in the lateral retraction of brown algae zygotes. Furthermore, it is interesting to note that in *Fucus,* M-M-rich alginates accumulate all around the fertilised spherical egg until the signal disappears when the zygote germinates (Torode et al., 2016). This ubiquitous localisation persists in the nearly spherical thallus cell of the two-celled *Fucus* embryo that constitutes the embryo proper (the rest of the two-celled embryo grows a rhizoid that degenerates). Therefore, the persistence of an anisotropic pattern of accumulation of M-M-rich alginates may be associated with lateral retraction of the zygote and limiting growth of the embryo in width.

In *Ectocarpus*, the location of alginates along the sporophytic filament does not correlate with cell differentiation, but with the amount of tensile stress at the surface of the cell, which is a combination of turgor, cell curvature and cell wall thickness (Rabillé et al., 2019b). Cell areas with lower cell wall thickness and a lower meridional curvature, which are two conditions of high tensile stress (Lockhart, 1965), have a stronger alginate signature compared with more curved areas with thicker cell walls (Rabillé et al., 2019). Furthermore, when probing the cell surface with atomic force microscopy, spherical cells with lower curvatures displayed a stiffer innermost cell wall layer than cylindrical cells with higher curvatures (Rabillé et al., 2019b). In the elongated *Saccharina* zygote, flanks have the lowest meridional and circumferential curvatures. Hence, because the thickness of the external cell wall seems to be similar throughout the egg and embryos (Fig. S5), the flanks likely experience the highest tensile stress. In this context, M-M-rich alginates positioned in the flanks of the zygote and embryo may prevent growth in this direction, and thereby promote the establishment of the apico-basal axis in Phase I embryos. In cell-walled organisms, there are several examples of envelopes that surround the embryo and maintain its mechanical balance. In *Arabidopsis*, a recent study reported a transient proteinaceous envelope around early globular-stage embryo (Harnvanichvech et al., 2023). It is thought to control the outward mechanical forces exerted by cell turgor in embryo cells, that escapes the control of the surrounding endosperm, which is deflating at that stage. However, in *Arabidopsis*, and in contrast to *Saccharina*, this control seems to be isotropic, the embryo being almost spherical at that stage.

In addition to the establishment of the embryo longitudinal axis through the retraction of the lateral sides of the fertilised egg, followed by the maintenance of the elongated shape by a corset of alginates as seen above, the production of cells almost twice as long as they are wide is the second major morphological anisotropy event. Remarkably, these cells are not oriented along the longest axis of the embryo, but perpendicular to it. They are produced by four synchronous cell divisions in the 4-cell stage embryo, parallel to the longest cell axis of their mother cell. The latter is already fairly anisotropic, with an LWR of 1.5. Therefore, these cell divisions diverge from the canonical rules of cell division according to which cells divide along the shortest path, i.e. perpendicularly to the longest cell axis (Besson and Dumais, 2011; Besson and Dumais, 2014; Errera, 1888; Hofmeister, 1863). In addition, the newly formed transverse cell walls seemed very dense in M-M-rich alginates, judging from the intensity of the signal generated by the BAM6 antibody. Cell divisions parallel to the main stress direction have been reported in plants (Louveaux et al., 2016; Rasmussen and Bellinger, 2018), where they are thought to prevent yielding in this direction. Therefore, in the *Saccharina* embryo, cell divisions perpendicular to the longitudinal axis may allow the embryo to better withstand circumferential tensile stress, which — in a prolate ellipsoid — is at its highest at the equatorial plane (Hejnowicz et al., 1977; Rabillé et al., 2019a).

In summary, at the zygote stage, the alginate corset may help maintain zygote elongation by preventing widening along the flanks. As the embryo grows, the stress increases and, in response to this stress, the cell division plane is established parallel to the circumferential stress, thereby reinforcing the mechanical resistance of the alginate corset. As a result, cells are highly anisotropic and incidentally further maintain the apico-basal axis of the embryo during Phase I, especially when the corset starts to disappear at the end of Phase I.

Although AFs are known to be involved in the formation of the cell wall in brown algae, and especially in the spatial arrangement of cellulose microfibrils (Karyophyllis et al., 2000; Katsaros et al., 2002), they have also been shown to be necessary for maintaining the strength of the cell wall in a Golgi -dependent manner (Bisgrove and Kropf, 2001). In *Saccharina*, treatment with the AF-depolymerising drug latrunculin B resulted in the loss of anisotropy of the shape of the zygote and the embryo, concomitantly with the loss of the anisotropic pattern of M-Mrich alginate accumulation (summarised in Fig. 6, bottom). This loss of anisotropy strongly suggests a causal relationship between three factors: cortical AFs (1) appear to control, probably together with yet uncharacterised protein complexes, the specific accumulation of M-M-rich alginates (2) along the flanks of the zygote, thereby stiffening the cell wall (3).

Interestingly, in contrast to what was observed in the external cell wall, the presence of mannuronate-rich alginates in the internal cell walls did not depend on the presence of AFs. AFs are necessary for cytokinesis of brown algal cells ((Mazéas et al., 2022; Nagasato et al., 2010)(Katsaros et al., 2013; Nagasato et al., 2010) and in our experiment, the cells labelled in Phase II *Saccharina* embryos had already divided. Therefore, depolymerisation of AFs by LatB would mainly affect the maintenance of the organisation of the previously formed cell wall. Although the signal corresponding to mannuronate-rich alginates seemed diffuse, potentially revealing some disorganisation of the cell wall, the signal remained homogeneously distributed at the surface of all internal cell walls. This diffuse pattern of distribution suggests that the maintenance of M-M-rich alginates in internal and external cell walls is controlled by different mechanisms. Nevertheless, the AFs organised mainly parallel to the Z-axis shows a potential relationship between AFs and mechanical feedback, reminiscent of cortical microtubules in plants (Hamant et al., 2019; Zhao et al., 2020).

In addition to the cortical AFs that control the formation of the corset, we have recently shown that the development of the longitudinal axis of the embryo depends on the connection with the maternal tissue. The integrity of the stalk, a structure corresponding to the remnantoogonium wall, is necessary for the zygote and embryo to develop the apico-basal axis (Boscq et al., 2024b). A signal diffusing from the stalk through the basal cell of the embryo appears to inhibit longitudinal divisions in the embryo (Boscq et al., 2024a). Upon embryo growth, the apical cells at a distance of at least 40 µm from the stalk can engage in longitudinal division. Beyond identifying the nature of this signal, the question of whether it controls the organisation of cortical AFs and the spatial pattern of M-M-rich alginates is the next important step in continuing to determine the mechanisms and signalling pathways that control *Sacharrina* embryogenesis. Before undertaking this endeavour, or perhaps concomitantly, it is necessary to confirm the role of this alginate corset in the elongation of the zygote, and thus in the establishment of the apico-basal axis of *Saccharina* embryo. Treatment with enzymes degrading the cell wall of brown algae, such as alginate lyases, results in a massive boost of bacterial contamination at the surface of the alga, due to the release of small glycans or monosaccharides in the medium. Therefore, a mechanical approach, or mutants with a reduced M-M-rich alginate composition are more promising to weaken the corset and test its effect on the shape anisotropy of the zygote and young embryo. However, as we have observed in this study, M-M-rich alginates are found mainly in the innermost layers of the external and internal cell walls, like M-G-rich alginates, and unlike G-G-rich alginates which are generally found in the outermost layers. Similar observations have been reported in *Ectocarpus* (Terauchi et al., 2016) and anticipated (McCully, 1965) in *Fucus*. This location suggests that mannuronates are delivered to the plasma membrane before being transformed into guluronates by the mannuronate C5-epimerases (Fischl et al., 2016) present in the cell walls (Mazéas et al., 2022). Alternatively, mannuronate-rich alginates may be degraded by alginate lyases recently identified in brown algae (Inoue and Ojima, 2019; Mazéas et al., 2022). These two hypotheses could explain the disappearance of the M-M-rich alginate corset at the end of Phase I, but on the whole they rule out the possibility of making an embryo without an M-M-rich alginate corset: first, M-M-rich alginates belong to the same cell wall layer as M-G alginates, which makes them difficult to destroy specifically, for example by laser irradiation; second, mutants lacking mannuronates will also lack guluronates, which will be highly detrimental to the alga, if not lethal.

In conclusion, here we established a functional relationship between the anisotropic growth of the embryo of *Saccharina latissima*, and the presence of a corset of mannuronate-rich alginates in the zygote, through the presence of cortical actin filaments. When AFs are depolymerised, the level of embryo anisotropy decreases and the corset disappears, replaced by a alginate layer homogeneously enveloping the zygote and embryo. This raises many questions, such as how cortical AFs located homogeneously around the zygote can control the formation of a corset only along the flanks of the embryo, how mannuronate alginates organised as a corset control the widening of the zygote and embryo, whereas they are organised as a layer distributed homogeneously in isotropic zygotes, which will need to be resolved by future work.

## Materials and methods

### Preparation of Saccharina embryos

Embryos were produced from fertilisation of female gametophytes as described in (Theodorou et al., 2021) and distributed in dishes with glass bottom (NEST). Briefly, gametogenesis was induced under white light conditions at 16 μmol photons m^-2^ s^-1^, 14:10 light:dark photoperiod and 13 °C and full Provasoli enriched seawater (PES) (Le Bail and Charrier, 2013; Theodorou et al., 2021). Then, the embryos were exposed to more intense white light (40 μmol photons m^-2^ s^-1^) to promote embryo growth.

### Embryo and cell morphometry

Data for embryo and cell morphometrics were taken from three sources: measurements of zygote dimensions (n=15) and previous measurements of control embryos (n=10) at stages ≥ 2-cells, published in (Boscq et al., 2024a) and (Boscq et al., 2024b). Zygote dimensions were measured on microscopic images using Fiji (Schindelin et al., 2012), and dimensions of embryos and their cells were computed by the software described in (Boscq et al., 2024a). For the embryo and cell measurements, the sliding window mean and the lowess function were computed using Python 3 (Van Rossum and Drake, 2023).

### BAM immunolabelling

BAM antibodies (Torode et al., 2015; Torode et al., 2016) were obtained from SeaProbes (Roscoff). Immunolocalisation was adapted from (Rabillé et al., 2019b). Briefly, embryos were fixed with 4% paraformaldehyde dissolved in H_2_O:sea water (NSW hereafter) (50:50) for 70 min at room temperature. Cells were washed once with NSW:PBS (50:50), twice with PBS for 10 min and blocked in PBS with 5% milk for 1 h at room temperature. The embryos were incubated overnight at 4°C with the BAM monoclonal primary antibodies (Torode et al., 2015; Torode et al., 2016) diluted 1:10 in PBS with 5% milk. Antibodies were washed with PBS three times for 5 min. The cells were then incubated overnight at 4°C with the secondary antibody anti-rat conjugated with FITC diluted 1:100 in PBS. Antibodies were washed three times for 10 min with PBS. The embryos were then stained for 30 min with 20 µM Calcofluor (Fluorescent Brightener 28, Sigma-Aldrich, St. Louis, MI, USA) and washed three times with PBS. Finally, the embryos were mounted in Vectashield (H-1000-10 (Vector®, with or without DAPI) and covered with a coverslip. Each labelling was repeated at least 3 times.

### Labelling actin filaments

The actin labelling protocol was optimised based on (Rabillé et al., 2018). Firstly, the material was incubated for 30 min in prefix solution containing 300 μM m-maleimido benzoic acid N-hydroxy succinimide ester (MBS; Sigma-Aldrich®), 0.2% Triton X-100 and 2% dimethyl sulphoxide (DMSO) in microtubule-stabilising buffer (MTB, 50 mM PIPES, 5 mM ethyleneglycol bis (aminoethyl ether)-tetraacetic acid (EGTA), 5 mM MgSO4.7H2O, 25 mM KCl, 4% NaCl, 2.5% polyvinylpyrrolidone 25 (PVP), 1 mM DL-dithiothreitol (DTT), pH7.4), in the dark and at room temperature. Fixation followed without washes, changing the medium to 2% PFA, 0.2% glutaraldehyde (GTA) and 2 U of rhodamine-conjugated phalloidin (Ph-Rh; R415, Invitrogen™) or Alexa Fluor™ 488-conjugated phalloidin (Ph-Alexa488; A12379, Invitrogen™) in MTB for 1.5 h, in the dark and at room temperature. Then the embryo tissue was washed with 1:1 MTB:PBS, and incubated in cell wall lysis buffer (2% w/v Cellulase Onozuka R-10 (Yakult Pharmaceutical industry Co., Ltd.), 2% w/v hemicellulase (Sigma-Aldrich®), 1% driselase (Sigma-Aldrich®), 1.5% macerozyme R-10 (Yakult Pharmaceutical industry Co., Ltd.), 50 U/ml alginate lyase-G (Station Biologique de Roscoff), 5 U Ph-Rh/ Ph-Alexa488 and 0.15% Triton X-100 in 1:1 MTB:PBS) for 10 min, in the dark and at room temperature. An extraction step followed a short series of washes. The tissues were incubated for 10 min in extraction solution (5% DMSO, 3% Triton X-100 and one of the phalloidin conjugates (2U) in PBS) in the dark and at room temperature. After washing with 1:1 MTB:PBS, the tissue was treated with the phallodin stain (15 U of the chosen phalloidin conjugate in 1:1 MTB:PBS) overnight at 4 °C in the dark. After washing, the nuclei were stained with DAPI. Vectashield H-1000-10 (Vector®) was used for mounting the samples.

The orientation of AFs was measured with the batch mode of FibrilTool included in the MT_Angle2Ablation workflow ((Boudaoud et al., 2014) https://github.com/VergerLab/FibrilTool_Batch_Workflow). The apico-basal axis of the embryo was set as reference (angle=0). The angles were processed with basic statistics in R.

### Latrunculin B treatments

Egg, zygotes and embryos of 2 to 8 cells were treated with a range of concentrations from 50 nM to 1 µM of latrunculin B (Sigma Aldrich) or incubated in 0.1% or 1% DMSO (Lat B solvant used as a control) for one week. Their growth was then monitored every day for up to 7 days under a bright field microscope (DMI8 inverted microscope, Leica Microsystems). Samples treated for 1 week with 1 µM LatB were used for the immunostaining experiments (results shown in Fig. S2).

### Image acquisition

Image data were acquired with a bright field and epifluorescence microscope DMI8 (Leica Microsystems) equipped with the colour camera DMC4500 (Leica Microsystems), Leica EL6000 compact light source and controlled with LAS X v3.0 (Leica Microsystems), and TCS SP5 AOBS inverted confocal microscope (Leica Microsystems) controlled by the LASAF v2.2.1 software (Leica Microsystems), with the objective HCX PL APO CS 20.0×0.70 DRY UV. The wavelengths of excitation of DAPI/Calcofluor white, FITC and chloroplasts were respectively 405 nm, 488 nm and 496 nm. The emission wavelength ranges were respectively 415-485 nm, 551-620 nm and 668-765 nm. The opening of the pinhole was 1AU. Line average and line accumulation were set to 1. Observation of rhodamine or Alexa Fluor^TM^ 488-phalloidin-labelled AFs was carried out using a TCS SP8 AOBS inverted confocal microscope (Leica Microsystems) with a 100X/N.A 1.4 objective.

The data were analysed using Fiji (Schindelin et al. 2012).

### Transmission electronic microscopy

Live embryos were left to grow for up to 6 days under white light, then fixed with 1% glutaraldehyde and 1% paraformaldehyde (both from Sigma-Aldrich) in sterile, 0.2 μm-filtered seawater for 2 h at 13°C. Then, the fixation medium was gradually changed to 0.1 M sodium cacodylate. Post-fixation consisted of incubating the thalli in 1% OsO4 in 0.1 M sodium cacodylate at 4°C overnight. Following washes with 0.1 M sodium cacodylate, the embryos were dehydrated using an ethanol:sodium cacodylate gradient. For infiltration, Spurr resin gradually replaced ethanol (Spurr, 1969), with fresh resin for the final step before polymerisation. Ultra-thin sections (50-70 nm) from the embedded embryos were mounted on copper grids (Formvar 400 mesh; Electron Microscopy Science©) and stained with 2% uranyl acetate for 10 min (Woods and Stirling, 2013) and 2% lead citrate for 3 min at room temperature (Reynolds, 1963). Sections were observed using a JEM-1400 Flash TEM microscope (JEOL Ltd.).

## Acknowledgements

S.B. was funded by the Bretagne Regional Council (“PRIMAXIS” project, grant number 1749) and Sorbonne University. I.T. was funded by an ARED grant from the Bretagne Regional Council (“PUZZLE” project) and NMBU. We are grateful to MITI CNRS for additional financial support and to Sophie Le Panse (Merimage platform, Roscoff Marine Station), for the TEM experiments. Part of the work was funded by the European Union (ERC, ALTER e-GROW, project number 101055148). Views and opinions expressed are those of the author(s) only and do not necessarily reflect those of the European Union or the European Research Council Executive Agency. Neither the European Union nor the granting authority can be held responsible for them. For purposes of open access dissemination, the authors applied for a CC BY-NC license for this document.

## Supplementary figures

**Fig. S1:**
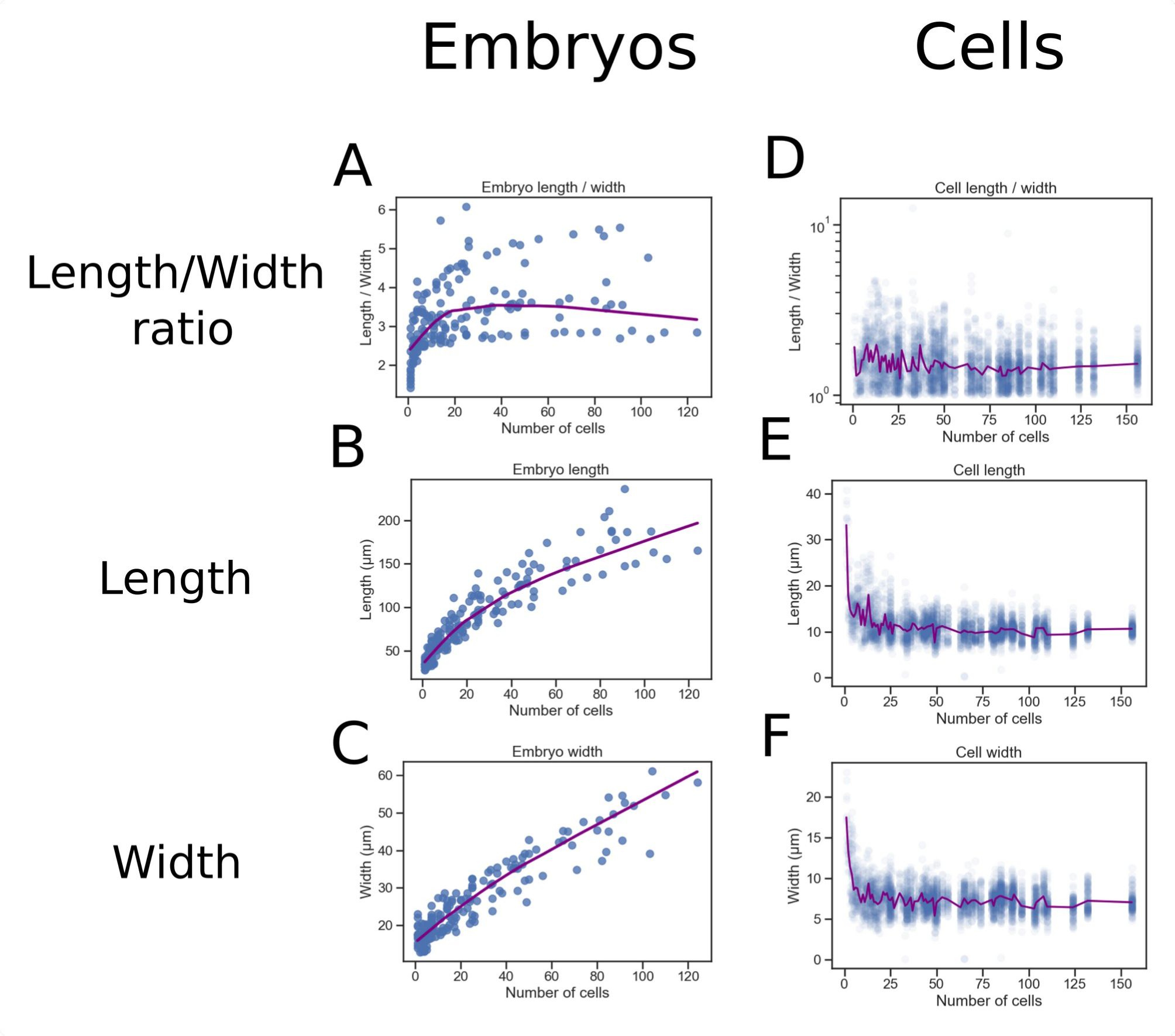
Cell size during *Saccharina* embryogenesis. Embryo and cell length/width ratio (A and D, respectively, µm), length (B and E, µm) and width (C and F, unit-less value) are plotted as a function of the developmental stage (expressed in the number of cells of the embryo), from the zygote stage to the 160-cell stage. (A,B,C): n = 17 embryos, the purple line shows the mean, smoothed using the lowess function; (D,E,F): Each blue circle represents the measurement of one cell. n = 17 embryos, with an average of eight cells per embryo. The purple line indicates the mean. Eggs (defined as single-celled organisms with an LWR ≤ 1.2) are excluded from these graphs.

**Figure S2.**
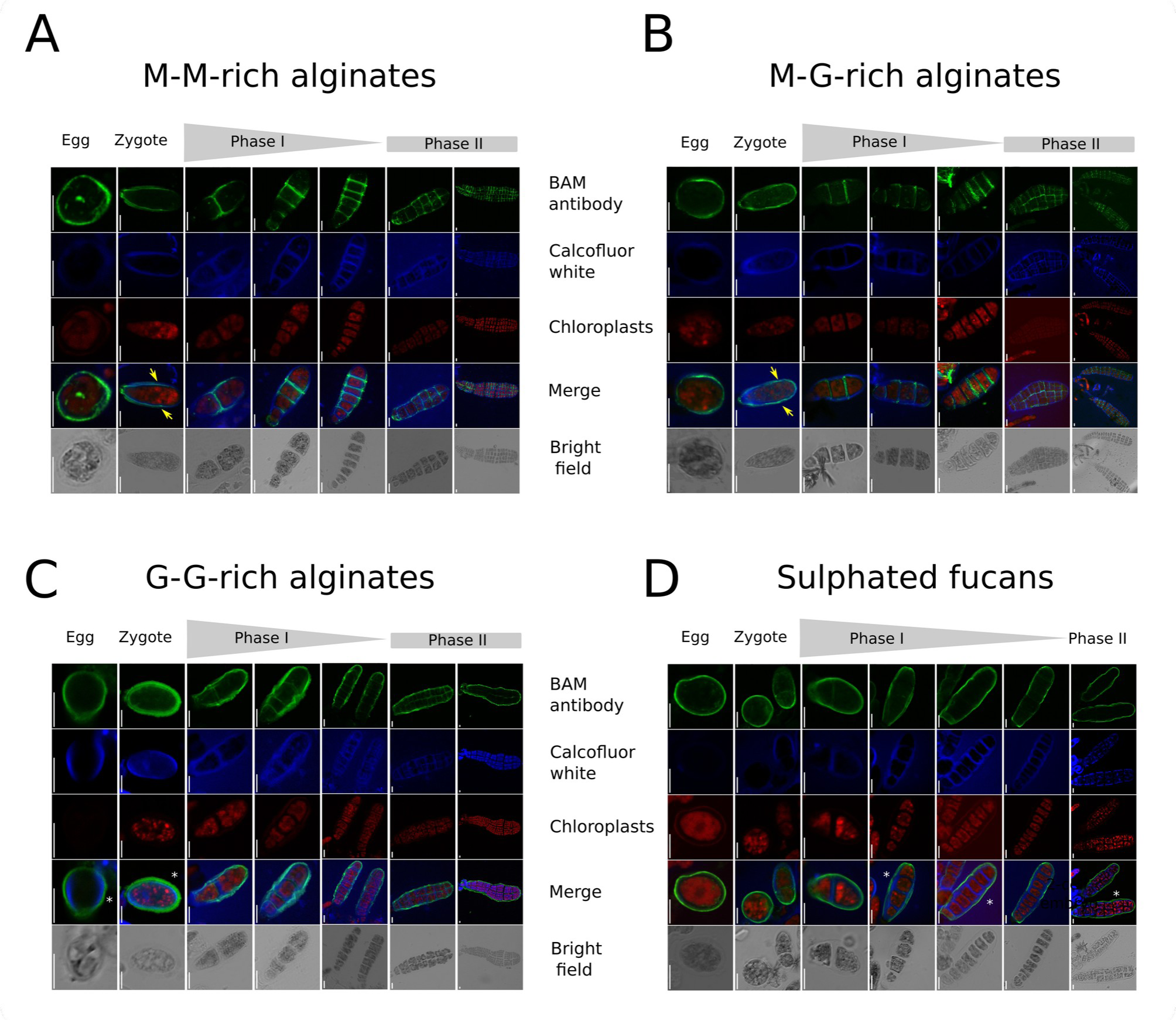
Eggs, zygotes and embryos labelled with BAM6, BAM7, BAM10 and BAM4 antibodies and Calcofluor white. BAM4-10 are detected with FITC-secondary antibodies (green, first row), cellulose with Calcofluor white (blue, second row). Red signal shows the autofluorescence of chloroplasts (third row). Merge of BAM, Calcofluor and autofluorescence is shown on the fourth row. Grey: bright field. (A) M-M-rich alginates labelled with BAM6; (B) M-G-rich alginates labelled with BAM7; (C) G-G-rich alginates labelled with BAM10; (D) sulphated fucans labelled with BAM4. White asterisks indicate where cellulose is observed more internally than G-G-rich alginates or sulphated fucans. Yellow arrows show where cellulose is more external than M-M- and M-G-rich alginates. Scale bars: 10 µm.

**Figure S3:**
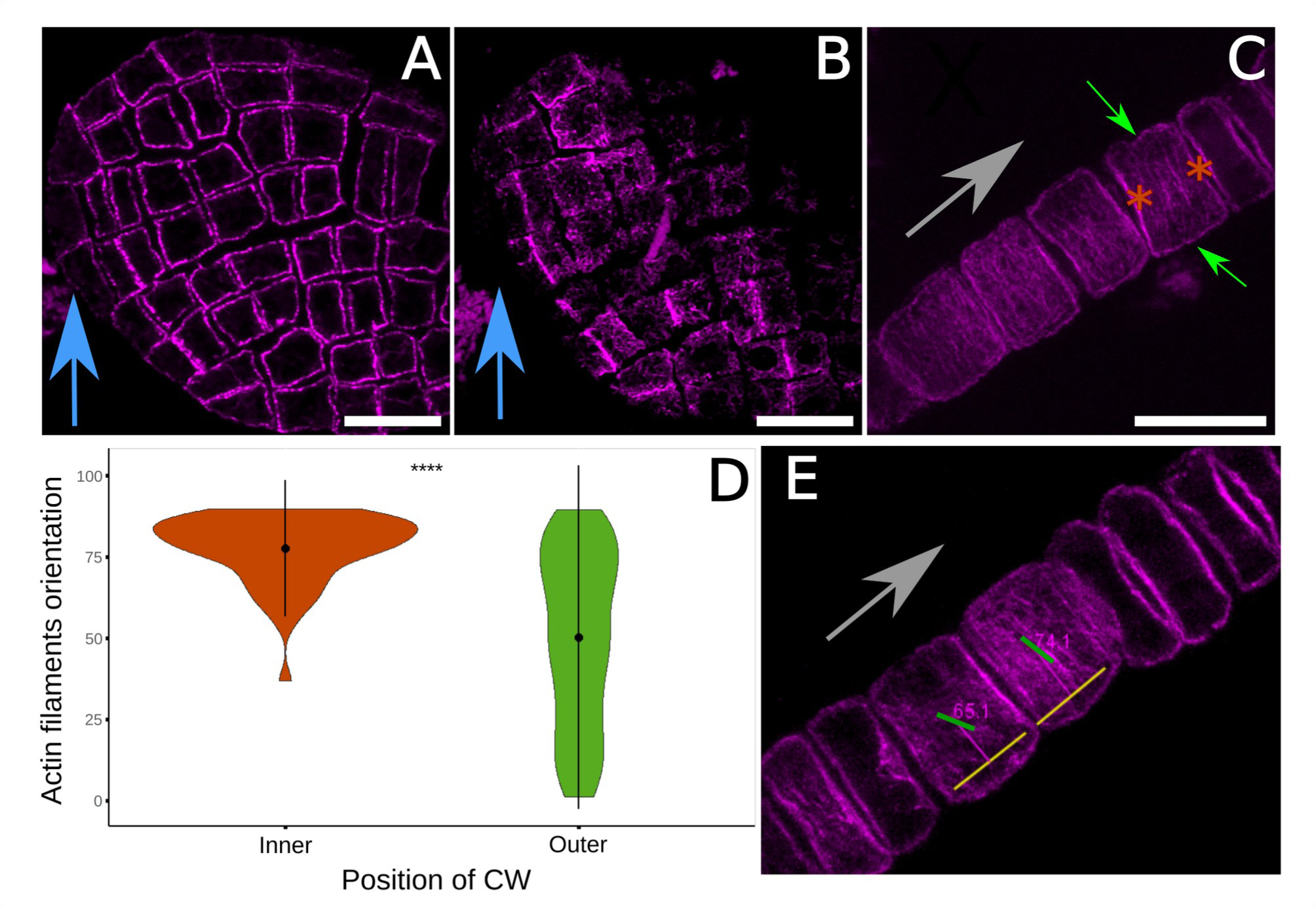
Anisotropic orientation of actin filaments during Phase II. Confocal images. (A) Median optical section of an apex from a Phase II embryo stained with rhodamine-phalloidin showing cortical actin filaments (AFs) (X-Y orientation). (B) Surface optical section of the same apex as in A (X-Y orientation) stained with rhodamine-phalloidin. (C) Surface longitudinal optical section (X-Z orientation): Cortical AFs from cell faces within the embryo lamina (owing to the longitudinal section, these are “internal faces”; brown asterisks) and from faces in contact with seawater (the “external” or outer faces, green arrows), stained with Alexa Fluor^TM^ 488-phalloidin. Note the different orientations of AFs compared with B. (D) Violin plots of AF orientation in same sections as in C, measured relative to the embryo apico-basal axis (X-axis; angle = 0) with MT_Angle2Ablation workflow in Fiji. **** = p-value < 0.0001. N = 54 cells for the cortical AFs of internal cell faces, observed along the Z-axis, and N = 315 cells for the outer cell faces in contact with seawater. (E) A visual example of the measurements. Yellow line: reference line, parallel to the apico-basal axis of the embryo (grey arrow); green line: the mean direction of the AFs, the mean angle is written in purple. The reference for surface optical section was the medio-lateral axis of the tissue (blue arrow). Bars: 15 μm.

**Figure S4.**
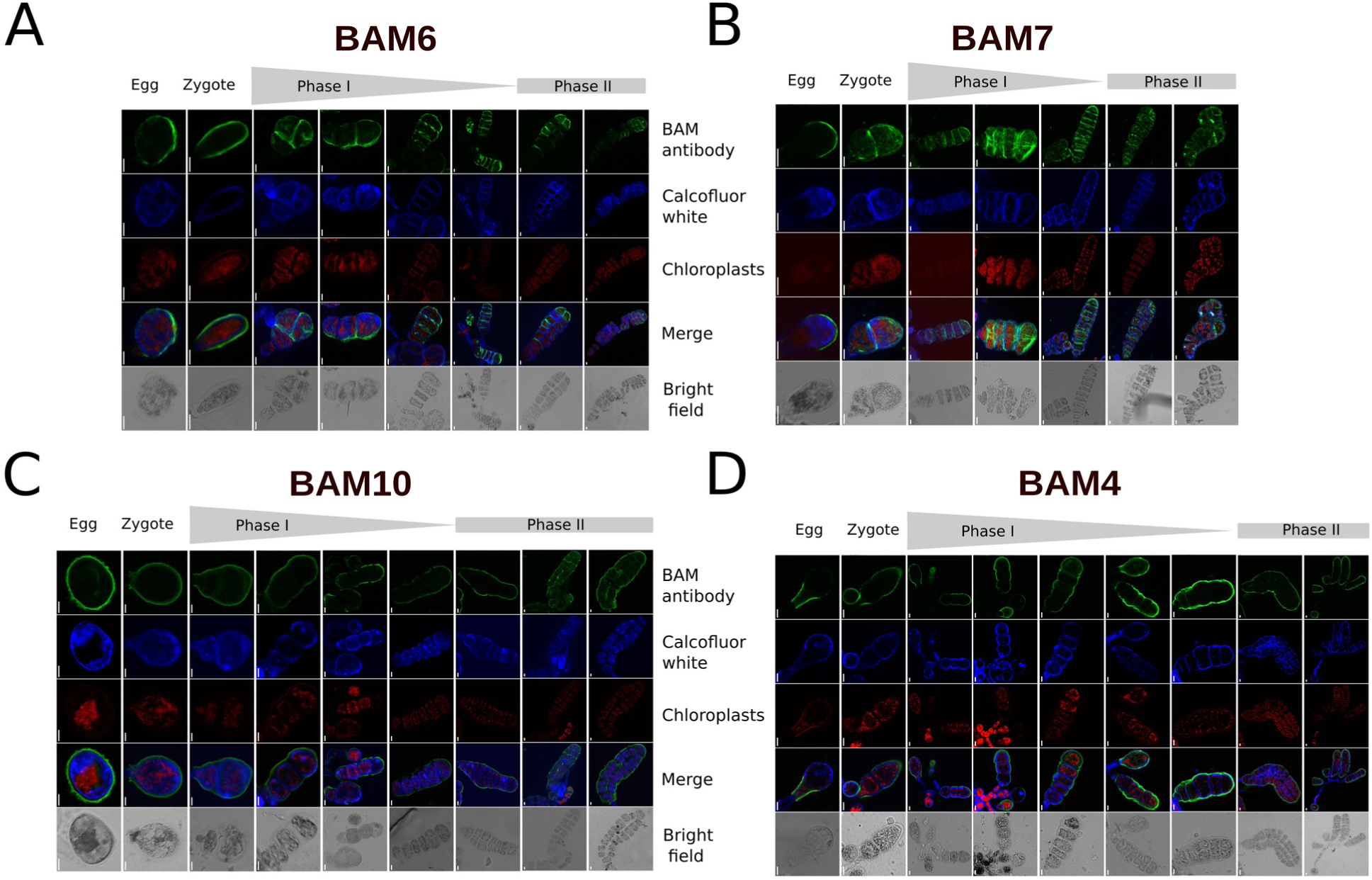
LatB-treated eggs, zygotes and embryos labelled with BAM6, BAM7, BAM10 and BAM4 antibodies and Calcofluor white. (A) M-M-rich alginates labelled with BAM6; (B) M-G-rich alginates labelled with BAM7; (C) G-G-rich alginates labelled with BAM10; (D) Sulphated fucans labelled with BAM4. Green: BAM antibodies (first row); Blue: Calcofluor white (cellulose, second row). Red signal shows the autofluorescence of chloroplasts (third row). Merge (fourth row); Grey: bright field (last row). Note that unfertilised eggs are not labelled because they have no cell wall at that stage. Scale bars: 10 µm.

**Figure S5.**
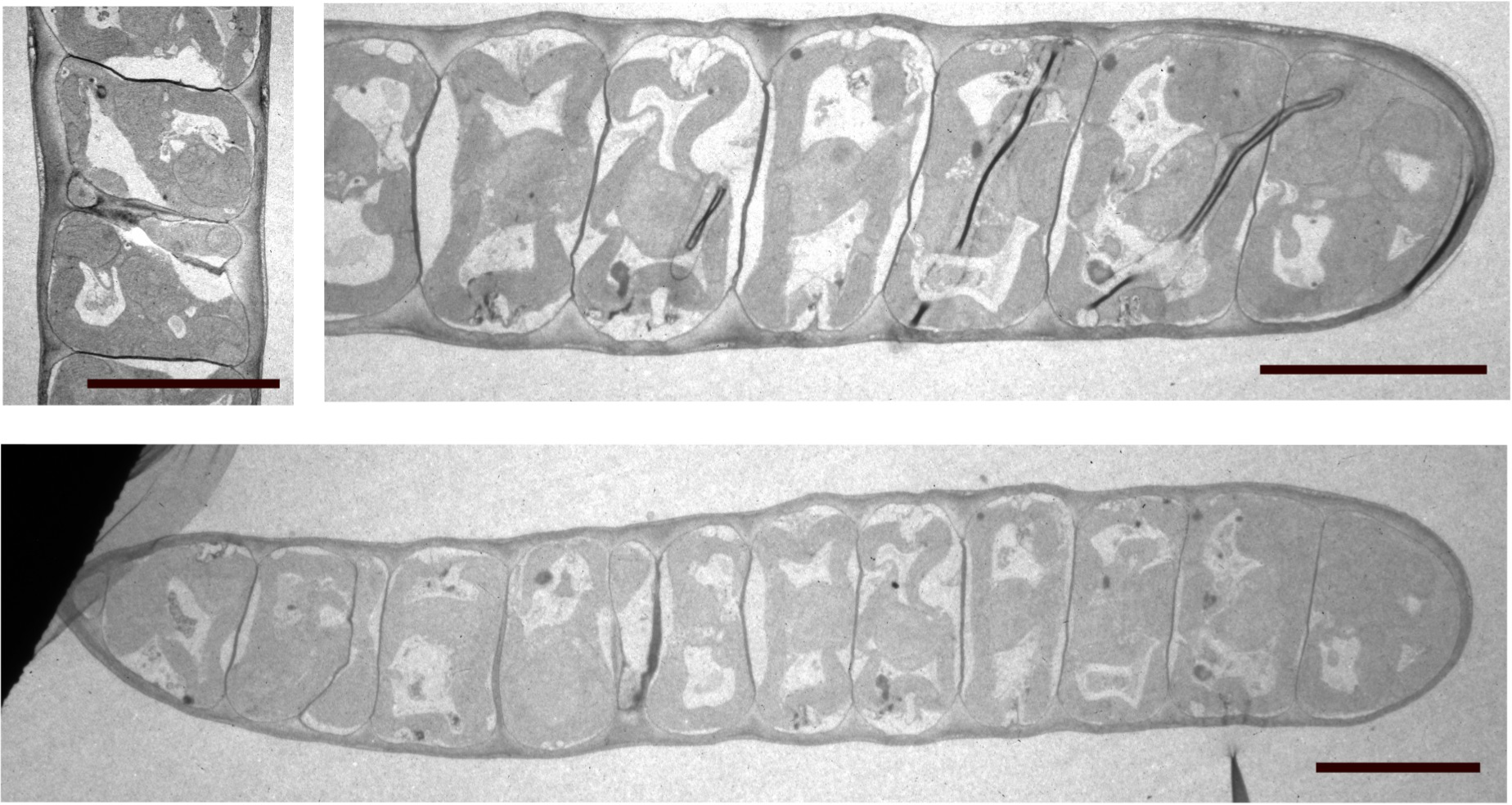
Cell wall thickness in *Saccharina* embryos. TEM observations of frontal or median sections of late Phase I (top) or early Phase II (bottom) embryos showing the thickness of the cell walls. Note the constant thickness of the outer cell wall along the embryo, and the different thicknesses between the outer cell wall and the transverse cell walls. Scale bar represents 10 µm.

**Table S1: Morphometrics of embryos and cells.** (A) Embryo (lamina) morphometrics; (B) Cell morphometrics

## Notes

### Competing Interest Statement

The authors have declared no competing interest.

